# Diminished cortical excitation and elevated inhibition during perceptual impairments in a mouse model of autism

**DOI:** 10.1101/657189

**Authors:** Joseph Del Rosario, Anderson Speed, Hayley Arrowood, Cara Motz, Machelle Pardue, Bilal Haider

## Abstract

Sensory impairments are a core feature of autism spectrum disorder (ASD). These impairments affect visual perception (Robertson and Baron-Cohen, 2017), and have been hypothesized to arise from imbalances in cortical excitatory and inhibitory activity (Rubenstein and Merzenich, 2003; Nelson and Valakh, 2015; Sohal and Rubenstein, 2019); however, there is little direct evidence testing this hypothesis in identified excitatory and inhibitory neurons during impairments of sensory perception. Several recent studies have examined cortical activity in transgenic mouse models of ASD (Goel et al., 2018; Antoine et al., 2019; Lazaro et al., 2019), but have provided conflicting evidence for excitatory versus inhibitory activity deficits. Here, we utilized a genetically relevant mouse model of ASD (CNTNAP2^−/−^ knockout, KO; Arking et al., 2008; Penagarikano et al., 2011) and directly recorded putative excitatory and inhibitory population spiking in primary visual cortex (V1) while measuring visual perceptual behavior (Speed et al., 2019). We found quantitative impairments in the speed, accuracy, and contrast sensitivity of visual perception in KO mice. These impairments were simultaneously associated with elevated inhibitory and diminished excitatory neuron activity evoked by visual stimuli during behavior, along with aberrant 3 – 10 Hz oscillations in superficial cortical layers 2/3 (L2/3). These results establish that perceptual deficits relevant for ASD can arise from diminished sensory activity of excitatory neurons in feedforward layers of cortical circuits.

## Introduction

Impaired sensory perception is a key feature of autism spectrum disorder (ASD) (Robertson and Baron-Cohen, 2017). Sensory disturbances may occur in >90% of individuals with ASD (Tavassoli et al., 2014), and are present early in development (Tomchek and Dunn, 2007). These sensory symptoms can predict later disease severity (Estes et al., 2015). Since there is detailed knowledge about the neural circuit basis of mammalian sensory processing, understanding impaired sensory perception in autism models provides an entry point for identifying neural circuit dysfunctions underlying core symptoms of ASD.

A prominent theory of ASD proposes that imbalanced excitatory-inhibitory activity ratios in cortex generate behavioral deficits (Rubenstein and Merzenich, 2003; Nelson and Valakh, 2015; Sohal and Rubenstein, 2019). However, little direct evidence for this hypothesis has been measured from identified excitatory and inhibitory neurons during quantifiable behavioral impairments. One recent study in a Fragile X syndrome mouse model of ASD found that reduction of inhibitory rather than excitatory activity in superficial cortical layers 2/3 (L2/3) correlated with sensory deficits (Goel et al., 2018). However, another study in the CNTNAP2^−/−^ KO mouse model of ASD found reduced and poorly coordinated excitatory activity in frontal cortex (Lazaro et al., 2019), but these excitatory activity deficits were not measured during sensory impairments or behavior. A third study suggests that these and multiple other ASD mouse models homeostatically compensate for deficits of excitatory and inhibitory activity, resulting in overall preserved sensory responsiveness (Antoine et al., 2019); crucially, this study also did not measure neural activity deficits during sensory perceptual impairments. It thus remains unresolved whether excitatory or inhibitory neural activity deficits best explain perceptual impairments in ASD mouse models.

There is extensive mechanistic knowledge about the excitatory and inhibitory basis of visual processing (Douglas and Martin, 2004; Isaacson and Scanziani, 2011; Priebe and Ferster, 2012), providing an ideal framework for resolving questions about neural activity deficits and perceptual impairments in ASD model mice. Remarkably, in individuals with ASD, deficits of visual processing arise as early as primary visual cortex (V1) (Robertson et al., 2014), but there remains considerable debate about how deficits in V1 activity might relate to perceptual impairments. We recently established that the state of activity in V1 plays a decisive role for trial-by-trial visual spatial perception in mice (Speed et al., 2019). This platform enabled us to here directly record putative excitatory and inhibitory neuron spiking in V1 of CNTNAP2^−/−^ KO mice while measuring the speed, accuracy, and contrast dependence of perceptual behavior. We found that KO mice showed multiple quantitative deficits in visual perception, and these were simultaneously associated with elevated inhibitory and reduced excitatory neuron activity along with aberrant low frequency network oscillations in the superficial layers of V1.

## Results

We trained both C57BL6J (wildtype, WT) and CNTNAP2^−/−^ knockout (KO) mice to report perception of spatially localized visual stimuli. Mice learned to lick for water rewards when visual stimuli (horizontally oriented Gabor gratings, see Methods) appeared on a screen (Fig. 1A). Stimuli appeared only after a mandatory period of no licking (0.5 – 6 s, randomized per trial), and rewards were delivered only upon the first lick during the stimulus response window (typically 1 – 1.5 s). We quantified perceptual performance using signal detection theory (Green and Swets, 1974). Our prior studies in WT mice showed that stimulus detection in the peripheral (monocular) visual field is more difficult than detection in the central (binocular) visual field (Speed et al., 2019). Here we found that KO mice had lower detection sensitivity than WT mice for stimuli appearing in these more difficult (monocular) spatial locations. Spatial detection sensitivity dropped to chance level at significantly nearer eccentricity in KO versus WT mice (49 ± 4°, n = 5 KO mice vs 72 ± 2°, n = 7 WT mice, *p<0.01* single tail rank sum). Since the goal of our study was to examine neural activity deficits in KO mice during repeatable and robust measurements of perceptual performance, here we focused on examining neural correlates of visual detection in the binocular visual field (central ±20°). This allowed us to measure large numbers of correct and incorrect behavioral trials, while presenting visual stimuli at the same spatial locations, visual contrasts, and durations for both WT and KO mice (see Methods).

**Figure 1.**
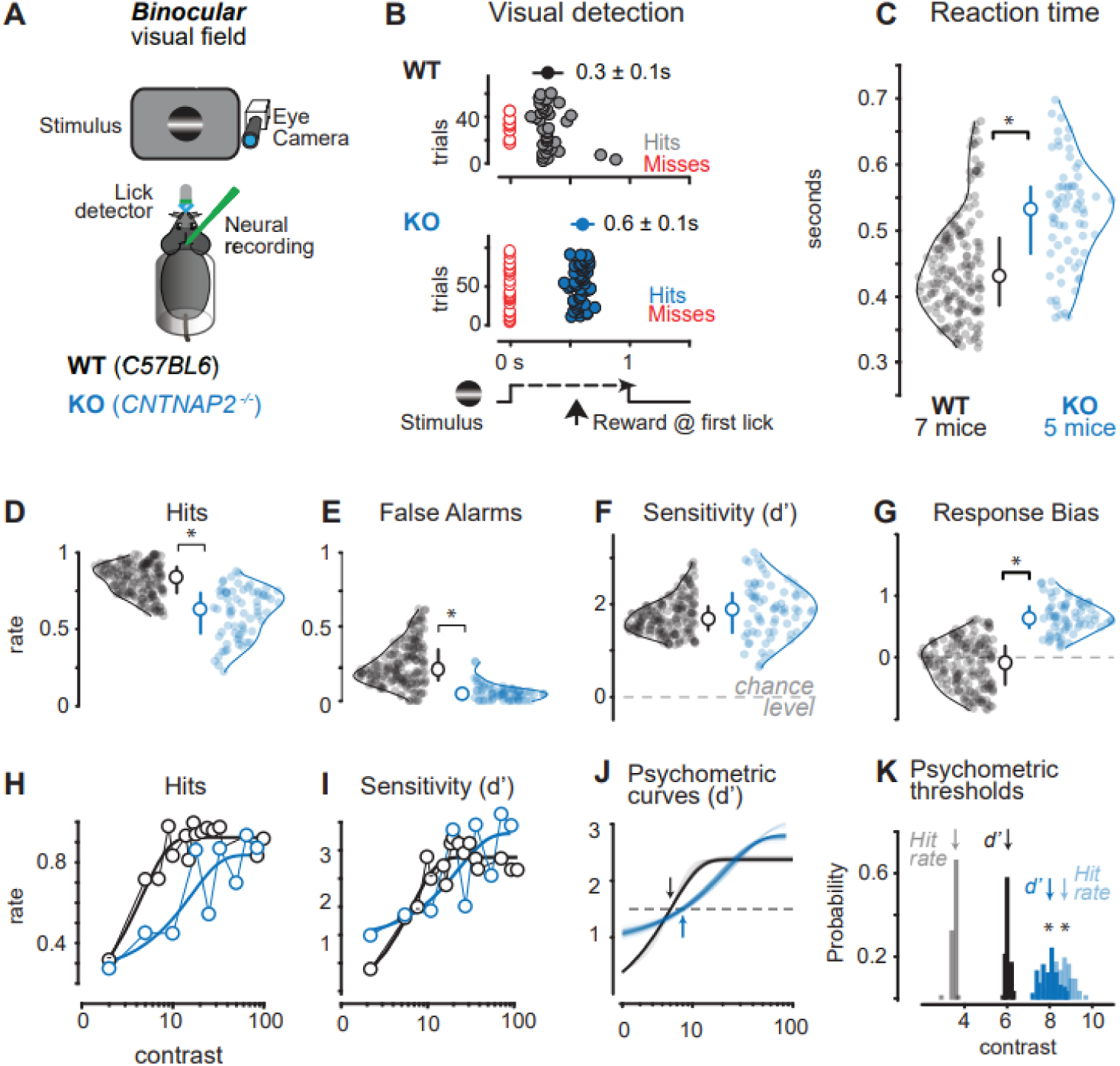
Impaired visual perceptual behavior in the CNTNAP2^−/−^ mouse model of ASD. **A.** Head-fixed mice reported detection of visual stimuli in the binocular visual field by licking. Pupil activity, neural activity, and licking was recorded simultaneously with behavior. C57BL6J (Wildtype, WT) in black, CNTNAP2^−/−^ (KO) in blue throughout. **B.** Example behavioral session shows detection latency (reaction time) was markedly slower for KO versus WT mice. Stimulus time course shown at bottom, with first lick times on correct trials (hits, colored circles) shown for individual consecutive trials (ordinate). Failures of detection (Misses) plotted in red. Average reaction times: WT, 0.3 ± 0.1s; KO, 0.6 ± 0.1s, mean ± SD reported throughout the figure. **C.** KO mice detected stimuli significantly more slowly than WT mice (KO: 0.52 ± 0.08, 71 sessions, 7 mice; WT: 0.45 ± 0.08, 187 sessions, 5 mice; *p < 0.01*, Wilcoxon rank sum throughout the figure). Average stimulus contrast was not significantly different across KO and WT mice (WT: 23 ± 24%; KO: 23 ± 22%). Circles show reaction time average per session. Median ± IQR plotted inside the distributions. **D.** KO mice showed significantly lower hit rates (KO: 0.6 ± 0.18; WT: 0.82 ± 0.12; *p < 0.01*). **E.** KO mice showed significantly lower false alarm rates (KO: 0.06 ± 0.07; WT: 0.24 ± 0.15; *p < 0.01*). **F.** Sensitivity index (d’) was not different between KO and WT mice (KO: 1.84 ± 0.71; WT: 1.74 ± 0.47; *p = 0.1*). **G.** KO mice showed higher criterion (c) indicating increased bias to withhold from responding (WT: 0.12 ± 0.43; KO: 0.65 ± 0.29; *p < 0.01*). Criterion was significantly greater than 0 for KO mice, but not for WT mice (WT: *p = 0.06*; KO: *p < 0.01*). C – G all during same behavioral trials and same mice. **H.** Hit rate as a function of contrast. Dark line is a psychometric fit (See Methods). **I.** Same as H, for d’. H-I during same sessions in same mice as C-G. **J.** Psychometric fit reliability from data in I (see Methods). Dashed line indicates d’ threshold (1.5), arrows indicate mean contrast values at threshold. Curves show 100 fits by resampling (see Methods). **K.** Contrast thresholds for hit rate (at 50% correct) and d’ (at 1.5) both significantly elevated in KO versus WT mice (hit rate contrast at threshold: WT=3.5 ± 0.1%, KO=8.7 ± 0.4%, *p<0.01*; d’ threshold: WT = 6.0 ± 0.1%, KO = 8.0 ± 0.4 %, *p<0.01*). Arrows indicated means of resampled psychometric curve threshold distributions (see Methods).

We first examined detection speed and accuracy for stimuli aggregated across all contrasts. KO mice detected binocular visual stimuli less frequently and more slowly than WT mice. KO mice showed significantly slower reaction times (Fig. 1B-C), andsignificantly fewer correct (Hit) trials of stimulus detection (Fig. 1D) than WT mice. However, KO mice also made fewer false alarms, which led to overall psychometric detection sensitivity (d’) that did not differ significantly from WT mice (Fig. 1E-F, see Methods). These measurements also revealed that KO mice held a significantly more conservative response bias than WTs (Fig. 1G). However, this conservative response bias was not simply explained by lower arousal—in fact, KO mice showed higher arousal (measured via pupil area) than WT mice before stimulus onset (Fig. 1 – figure supplement 1), and both WT and KO mice showed relatively lower arousal preceding correct detection, consistent with prior reports (McGinley et al., 2015; Speed et al., 2019). Higher arousal did not lead to higher distractibility in KO mice. We measured the rate of premature responses (stray licks) during pre-stimulus periods as a proxy of distraction (impulse control; Fonseca et al., 2015). KO mice showed significantly fewer premature responses(1.41 ± 0.16 stray licks / trial) than WT mice (2.42 ± 0.11, *p < 0.01*, rank sum test); moreover, premature responses in KO mice were not significantly modulated by arousal (1.5 ± 0.1 stray licks / trial versus 1.6 ± 0.1, sorted relative to median pupil area, *p = 0.73*, rank sum test). This suggests that KO mice do not show greater distractibility as a function of arousal. In addition, response vigor was comparable for KO and WT mice on Hit trials (similar licking frequencies, one fewer lick per reward in KO mice, Fig. 1 – figure supplement 1). Overall, these measurements argue against gross motor deficits or arousal differences as main factors underlying behavioral impairments.

We next found clear differences in detection performance as a function of stimulus contrast. KO mice showed far lower hit rates for stimuli of low and medium contrast as compared to WT mice (Fig. 1H), with little difference for high contrast stimuli. Accordingly, psychometric detection sensitivity (d’) as a function of contrast was both reduced and less steep in KO mice (Fig. 1I). We next fit psychometric functions (for both hit rates and d’) using a resampling approach across subsets of the data (Busse et al., 2011), and found KO mice showed significantly shallower slope of the psychometric function (Fig.1 – figure supplement 2), and elevated threshold contrast (Fig. 1K; threshold at d’ ≥ 1.5). The weaker behavioral sensitivity to contrast in KO mice was also evident in reaction times during these same trials (Fig.1 – figure supplement 2). Taken together these results show that KO mice have impaired psychometric sensitivity when considering detection across a range of stimulus contrast.

We then measured visual responses in primary visual cortex (V1), and observed alterations in visually-evoked excitatory and inhibitory firing, but only during wakefulness. We measured fast spiking (FS, putative inhibitory) and regular spiking (RS, putative excitatory) neuron populations (Fig. 2, Fig. 2 – figure supplement 1) with silicon probe recordings across layers of V1. During anesthesia, we observed no differences between KO and WT mice in the overall distributions of visually-evoked spiking in either RS or FS neurons (Fig. 2C, D). However, recordings during wakefulness (but in the absence of the behavioral task) revealed that visually-evoked spiking of FS neurons in KO mice was significantly elevated versus WTs, while RS neuron responses were significantly reduced during the same recordings (Fig. 2E-H).

**Figure 2.**
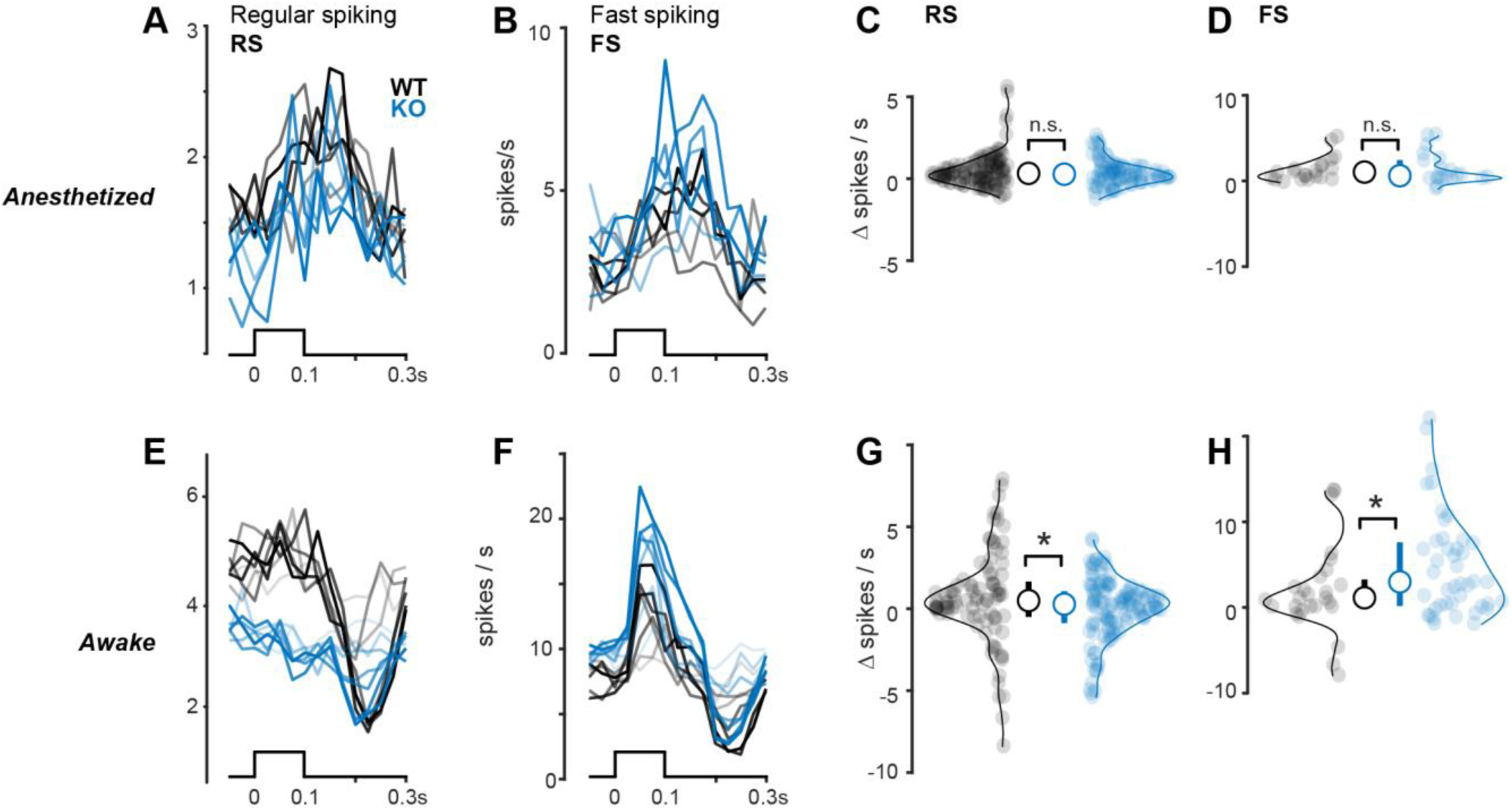
Reduced excitatory and elevated inhibitory neuron activity depends upon brain state. **A.** Regular spiking (RS) putative excitatory neuron firing to black or white bars (bottom) presented from 5-100% contrast during anesthesia. Stimulus contrast level indicated by line transparency. Spikes binned at 25 ms. **B.** Same as A, for fast spiking (FS) putative inhibitory neurons. **C.** No significant difference between stimulus-evoked activity in RS neurons during anesthesia (WT: 0.51 ± 0.12 spikes per s, n = 129 neurons; KO: 0.32 ± 0.09, n = 96 neurons, *p = 0.27*, 1-tail Wilcoxon rank sum test throughout the figure). **D.** Same as C for FS neurons (WT: 1.44 ± 0.44 spikes per s, n = 23 neurons; KO: 1.43 ± 0.45, n = 22 neurons, *p = 0.49*). **E.** Same as A, during wakefulness. **F.** Same as B, during wakefulness. **G.** Same as C, during wakefulness. RS neuron responses are significantly reduced in KO mice during wakefulness (WT: 0.49 ± 0.37, n = 95 neurons; KO: 0.03 ± 0.25, n = 129 neurons, *p<0.05*) **H.** Same as D, during wakefulness. FS neuron responses are significantly elevated in KO mice during wakefulness (WT: 1.90 ± 1.19, n = 45 neurons; KO: 5.10 ± 1.04, n = 29 neurons, *p<0.05*)

Consistent with behavioral responses, neural responses in awake KO mice also showed aberrant contrast dependence. We plotted spiking as a function of contrast (spaced from 5 — 100% contrast; Fig. 2 – figure supplement 2) and found that the FS neurons in KO mice had steeper contrast dependence than WT mice (linear fits, slope of firing rate versus contrast: KO, 6.1± 0.3; WT, 4.3 ± 0.3; *p<0.01*; Wilcoxon rank sum; mean ± SD); during these same experiments, contrast dependence of RS neuron firing in KO mice was significantly weaker than WT mice (slope of firing rate versus contrast: WT, 0.5 ± 0.05; KO: −0.21 ± 0.02; *p<0.01*; Wilcoxon rank sum; mean ± SD).

Aberrant neural responses in KO mice were not explained by differences in baseline excitability, receptive fields, response latencies, or retinal responses. First, baseline firing rates in the absence of visual stimuli were not different in awake WT and KO mice (RS: WT = 2.55 ± 0.32; KO = 2.96 ± 0.32; *p=0.39*; FS: WT = 6.16 ± 1.35; KO = 6.53 ± 1.14; *p=0.43*, single-tail rank sum test). Second, receptive field properties (spatial extent and width) were comparable in WT and KO mice (Fig. 2 – figure supplement 1). Third, the amplitude and latency of LFP responses during wakefulness was in fact significantly faster in KO mice across all layers (Fig. 2 – figure supplement 2). Lastly, perceptual impairments were not explainable by slower or diminished visual responses in the retina—these were nearly identical in KO and WT mice across a wide range of light intensities (Fig. 2 – figure supplement 2).

During perceptual behavior, KO mice showed elevated FS activity and reduced RS activity. Recordings in binocular V1 (Fig. 2 – figure supplement 1) revealed an overall greater number of visually-evoked spikes in FS neurons in KO versus WT mice on correct detection (Hit) trials at perceptual threshold (Fig. 3B), and simultaneously fewer visually-evoked spikes in RS neurons (Fig. 3F). These effects were observed in the first 0.2s following stimulus onset, but were evident even when including spikes up to 0.3s after stimulus onset (not shown). Again, these changes in KO mice during behavior were not due to lower arousal: pupil was larger in KO versus WT mice (Fig. 1 – figure supplement 1) and pupil dilation in KO mice resulted in significantly higher (not lower) firing rates (Fig. 3 – figure supplement 1, r = 0.98, *p < 0.01*). Moreover, these differences were not accompanied by gross alterations in the timing of excitatory and inhibitory firing in KO mice (Fig. 3-figure supplement 2). During behavior, the relationship between stimulus contrast and firing rate was significantly shallower in KO versus WT mice, in both FS (contrast vs firing rate linear fit slopes: WT, 5.6 ± 0.20; KO, 3.6 ± 0.01, *p < 0.01*, Wilcoxon rank sum test) and RS neurons (linear fit slopes: WT, 1.8 ± 0.04; KO, 0.1 ± 0.01, p < 0.01 Wilcoxon rank sum test). Overall, the weakest relationship between stimulus contrast and firing rate was in RS neurons of KO mice (*p<0.01*, Kruskal-Wallis test followed by multiple comparisons). These deficits in RS neurons are consistent with weaker contrast responses in awake mice outside of the task (Fig. 2-figure supplement 2), even though stimulus contrast range during behavior was far lower (5 – 35% contrast).

**Figure 3.**
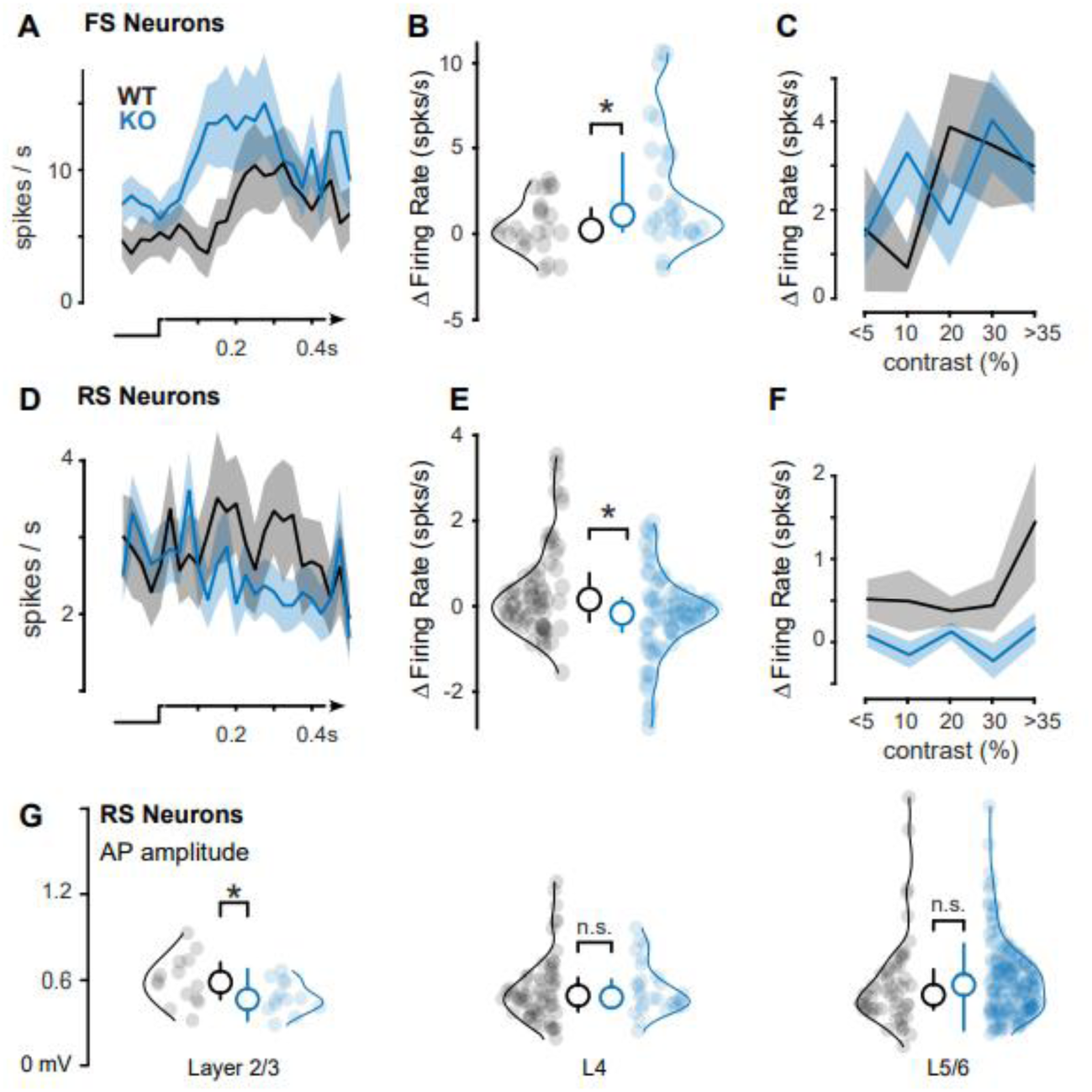
Enhanced FS and diminished RS visual responses at discrimination threshold. **A.** Peristimulus time histograms of FS neurons during perceptual detection of visual stimuli (bottom) at contrasts defined by the discrimination threshold (~5% for WT mice, ~10% for KO mice, see Fig. 1I-K). **B.** KO mice have enhanced FS responses to visual stimuli during perceptual detection (WT: 1.58 ± 1.27 spikes per s, n = 20 neurons; ASD: 3.29 ± 1.15, n = 38 neurons, *p<0.05*, 1-tail Wilcoxon rank sum test; Δfiring rate calculated as difference from pre-stimulus baseline, see Methods) **C.** FS neuron responses as a function of contrast (binned; see Methods). **D.** Same as A, for RS neurons **E.** Same as B, for RS neurons. KO mice have diminished RS responses during perceptual detection (WT: 0.52 ± 0.21 spikes per s, n = 49 neurons; ASD: −0.15 ± 0.18, n = 103 neurons, *p<0.01*, 1-tail Wilcoxon rank sum test) **F.** Same as C, for RS neurons. **G.** Action potential (AP) amplitudes significantly smaller in L2/3 RS neurons in KO mice (0.48 ± 0.03 mV; mean ± SEM n = 13; 0.59 ± 0.04 mV, n = 14; *p<0.05*, 1-tail Wilcoxon rank sum test). No differences in L4 (KO: 0.54 ± 0.03; n = 28; WT: 0.54 ± 0.03, n = 59; *p=0.35*) or L5/6 (KO: 0.59 ± 0.02; n = 178; WT: 0.58 ± 0.04, n = 53; *p=0.12*). Neurons aggregated across awake recordings. Median ± IQR plotted inside distributions.

Deficits in RS neuron activity in KO mice were most pronounced in Layer 2/3 (L2/3). We first examined spiking responses across layers (identified using current source density analysis; Fig. 2 – figure supplement1), and observed that we isolated far fewer neurons in L2/3 of KO versus WT mice. We aggregated across all awake recordings (both inside and outside of the task) and found that L2/3 excitatory neurons in KO versus WT mice showed significantly smaller action potential waveform amplitudes (Fig. 3G), consistent with prior reports (Scott et al., 2019). Importantly, in these same recording sessions, RS and FS neurons in L4 and L5/6 did not show significant differences in action potential amplitudes or activity profiles (L4: FS, *p=0.27*; RS, *p = 0.35*; L5/6: FS, *p=0.29*, RS, *p = 0.12*; rank sum tests), arguing against a global AP deficit in KO mice. We performed several further control measures to assess if differences in L2/3 RS activity were due to experimental conditions in KO mice. First, unresolvable background activity levels were comparable (or smaller) in KO versus WT mice, suggesting that reduced single-unit isolation in L2/3 was not due to higher background levels (Fig. 3 – figure supplement 3). Second, AP amplitudes were orders of magnitude larger than the background activity levels in both WT and KO recordings (Fig. 3-figure supplement 3), indicating that recording signal quality was not a factor. Third, the overall percentage of L2/3 RS neurons within a recording was not significantly different in WT and KO mice (WT: 4.2 ± 3.0%, n = 15 recordings; KO: 0.5 ± 0.4%, n = 24 recordings; mean ± SEM, *p=0.13*, one-tail rank sum test). Fourth, the percentage of isolated RS neurons did not decrease uniformly across all layers of KO mice (L5/6 – WT: 61 ± 7%, KO: 83 ± 6%, *p<0.05*, one-tail rank sum test; L4 – WT: 35 ± 7%, KO: 17 ± 6%, *p<0.05*, one-tail rank sum test). Lastly, the majority of L2/3 RS neurons in both WT and KO mice were isolated across multiple consecutive recording sessions spanning 1-2 days after the initial craniotomy (WT, 66.7% of total isolated after initial craniotomy; KO, 51% of total), suggesting that the quality and longevity of experimental preparations were comparable in WT and KO mice.

We next examined visually-evoked LFP responses, and these were also most strongly reduced in L2/3. We first measured responses in awake, non-task conditions (Fig. 4A), and found that LFP responses in L2/3 were significantly reduced at all contrasts compared to responses in WT mice. We then measured the difference in LFP response amplitude between WT and KO mice within layers, using the range of contrasts presented during the behavioral task, (5 – 35%), and found the largest reductions in L2/3 (*p<0.01*, repeated measures ANOVA followed by multiple comparisons, significant effect of layer).

**Figure 4.**
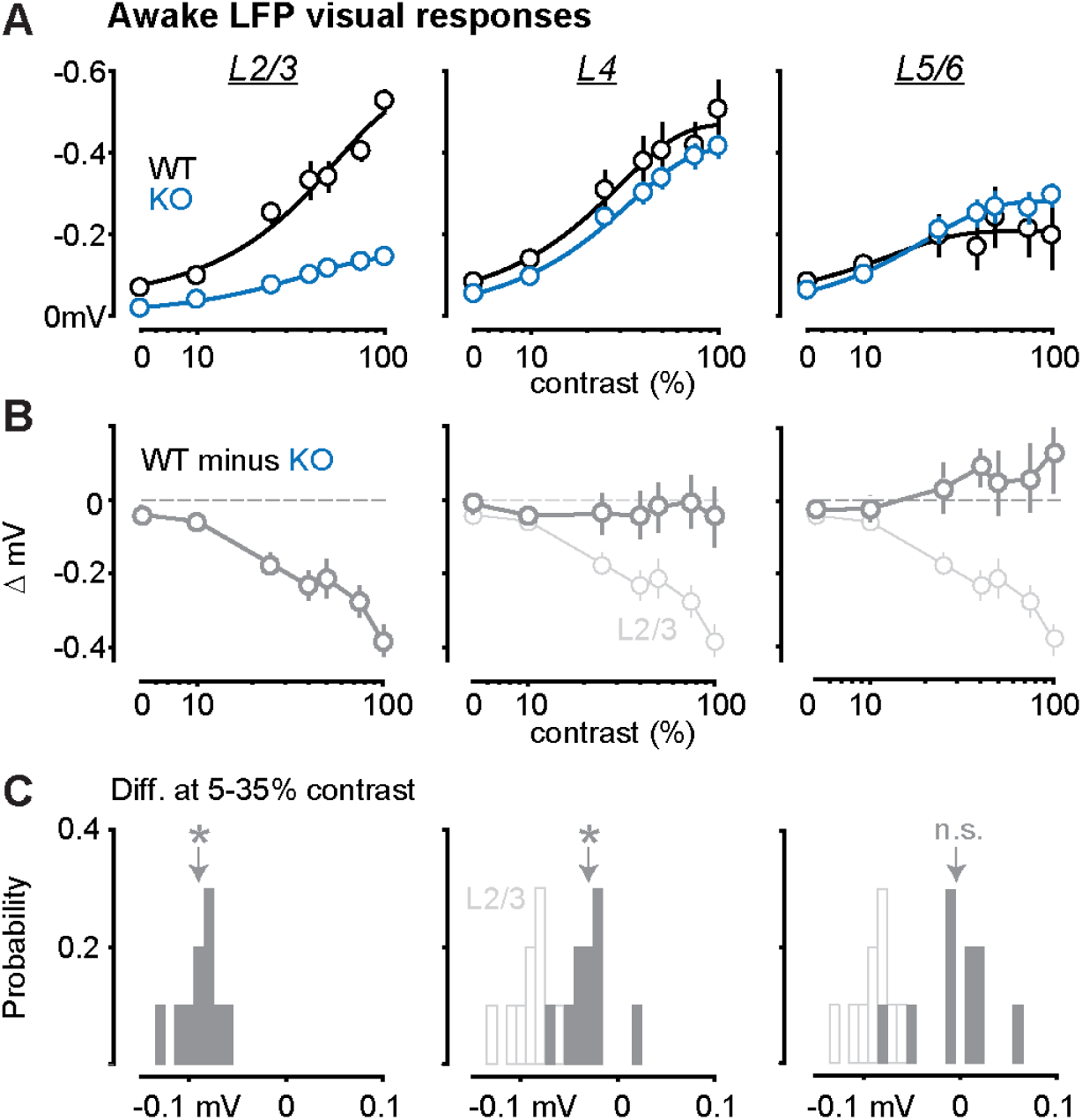
Strong reduction of LFP responses in L2/3 of KO mice. **A.** Contrast dependence of awake LFP responses, split by layers (L2/3, L4, L5/6; left to right). Ordinate reversed for visualization. Dark lines are Weibull fits (see Methods). **B.** Difference between WT and KO LFP contrast responses. L2/3 (left, grey), has the greatest reduction in stimulus evoked LFP compared to L4 (middle), and L5/6 (right) L2/3: −0.19 ± 0.014 mV, *p<0.01*; L4: −0.029 ± 0.007 mV, *p=0.23*; L5/6: 0.046 ± 0.011 mV, *p<0.01*, repeated measurements ANOVA followed by multiple comparisons, significant effect of layer. **C.** Probability distributions of LFP amplitude differences (see Methods). L2/3 responses (left) in KO mice are significantly reduced (−0.09 ± 0.002 mV, *p<0.01*, signed rank). L4 responses are slightly, but significantly reduced (−0.03 ± 0.003 mV, *p<0.05*). L5/6 are not different (−0.004 ± 0.004 mV, *p=0.77*). Distributions calculated from contrasts matching behavioral contrast range (2-35%). Open histograms replot L2/3 distribution for comparison to L4 and L5/6.

LFP responses in L2/3 of KO mice were also significantly reduced during perceptual behavior. On both correct and incorrect detection trials at psychometric threshold contrasts, L2/3 LFP responses to gratings were significantly smaller than those in WT mice (Fig. 5A). These results were not confounded by slower neural response latencies: there were no significant differences in LFP response latency to gratings successfully detected for rewards in WT versus KO mice (L2/3: WT=78 ± 4 ms, KO=75 ± 3 ms, *p=0.65;* L4: WT=78 ± 4 ms, KO=76 ± 3 ms; *p=0.8*; L5/6: WT=71 ± 3 ms, KO=70 ± 2 ms; *p=0.7)*. If anything, KO neural responses were slightly faster than WTs, consistent with passive awake recordings (Fig. 2 – figure supplement 2), and prior findings (Bertone et al., 2005). These weaker LFP responses in L2/3 of KO mice were apparent despite psychometric thresholds at higher stimulus contrast than WT mice.

**Figure 5.**
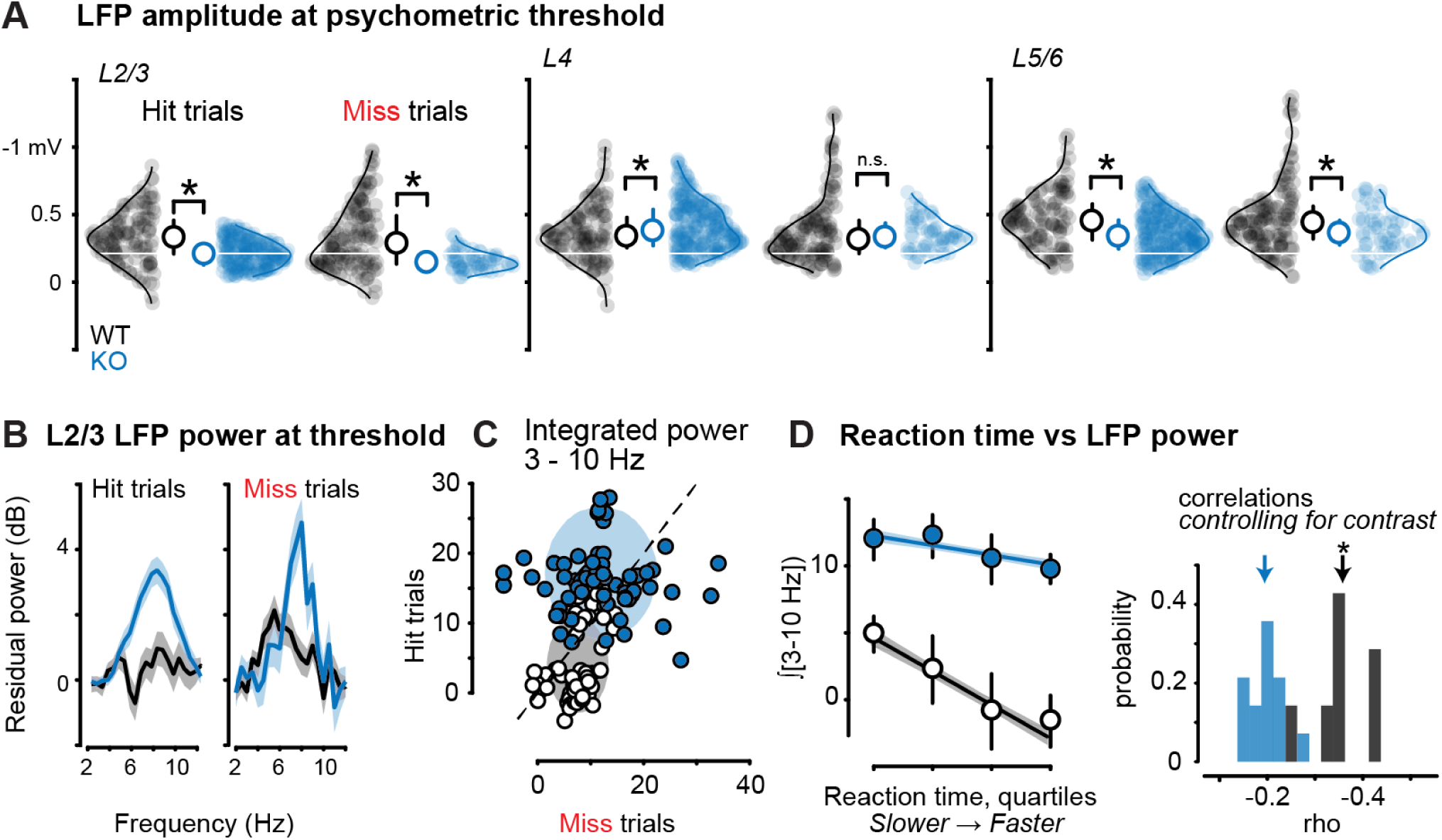
Aberrant neural activity in L2/3 correlates with perceptual impairments in KO mice. **A.** L2/3 stimulus evoked LFP amplitudes at discrimination threshold (see 1J-K) are reduced on both correct and incorrect trials (WT Hits: −0.33 ± 0.017 mV, n = 533 trials, 7 recordings in 3 mice; KO Hits: −0.21 ± 0.007 mV, n = 1402 trials, 15 recordings in 3 mice, p<0.01 Wilcoxon rank sum; WT Misses: −0.33 ± 0.020 mV, n = 211 trials; KO Misses = −0.16 ± 0.009 mV, n = 423 trials, *p<0.01*). Middle, L4 responses on Hit (WT: −0.36 ± 0.017 mV; KO = −0.41 ± 0.011, *p<0.05*) and Miss trials (WT:−0.38 ± 0.020 mV; KO: −0.35 ± 0.016 mV, *p=0.52*). Right, L5/6 responses on Hit (WT-0.46 ± 0.018 mV, KO = −0.36 ± 0.010 mV, *p<0.01*) and miss trials (WT: −0.48 ± 0.021.0, KO = −0.38 ± 0.021, *p<0.05*). L4 and L5/6 responses during detection are not reduced to the same magnitude as L2/3 in KO mice (L2/3 median indicated by white horizontal line). **B.** Significantly elevated low frequency (3 – 10 Hz) LFP power in L2/3 of KO versus WT mice on both Hit trials (KO: 13.27 ± 1.55; WT: 2.45 ± 1.78; *p<0.01*) and Miss trials (KO: 11.98 ± 2.51; WT: 7.03 ± 1.57; *p<0.05*). Mean ± SEM of integrated power 3 – 10 Hz at psychometric threshold. See Methods for residual power calculation. **C.** Integrated 3-10 Hz residual power was greater on Misses vs. Hits in WT mice (Hits: 4.01 ± 0.69; Misses: 7.56 ± 0.48, *p<0.01*, sign rank), but not in KO mice (Hits: 17.22 ± 0.76; Misses: 11.95 ± 0.93, *p<0.01*, sign rank). Shaded regions show 2-D Gaussian fit (± 1σ). Power calculated per channel recorded in L2/3 (WT: n = 58, KO: 68). **D.** Left, Low frequency LFP power was significantly and negatively correlated with reaction time in WT mice (linear regression model: 48 ± 12% variance explained within mouse, p<0.05 r^2^ = 0.35; *p<0.05*), but not in KO mice (31 ± 8% within mouse, *p=0.22*, r^2^ = 0.19; p=0.08). Single trial reaction times (Hit trials) were binned into quartiles, and single trial integrated 3-10 Hz LFP power was averaged within quartile for all WT and KO trials (mean ± SEM). Shaded regions are bootstrap error of fits (see Methods). Right, correlations between low frequency LFP power and reaction time are not explained by contrast. Partial correlations between the L2/3 power and reaction time, accounting for contrast, were significantly higher in WT mice (WT: 0.36 ± 0.06, p<0.01; ASD 0.19 ± 0.04, *p=0.08*; see Methods).

During perceptual behavior, L2/3 LFP responses in KO mice displayed aberrant low frequency power, and this degraded the relationship of neural activity to perceptual performance. Remarkably, 3 – 10 Hz LFP power evoked by stimuli at psychometric threshold contrast in MO mice was significantly elevated on both correct and incorrect detection trials (Fig. 5B), whereas in WT mice elevated 3-10 Hz LFP power was evoked preferentially on Miss trials, consistent with several prior studies (Einstein et al., 2017; McBride et al., 2019; Speed et al., 2019). When examined within recordings and within behavioral sessions, low frequency LFP power in L2/3 was significantly greater in KO mice on Hit rather than Miss trials (Fig. 5B; Integrated 3 - 10 Hz power in KO mice: Hits, 17.2 ± 0.8; Misses, 12.0 ± 0.9, *p<0.01* Wilcoxon signed rank test; in WT mice: 4.0 ± 0.7; Misses, 7.6 ± 0.5, *p<0.01*, Wilcoxon signed rank test; mean ± SEM), and provided poorer decoding of perceptual outcome at threshold (AUROC for Hits vs. Misses: WT=0.541 ± 0.003, mean +/− SEM, KO=0.525 ± 0.002, *p<0.01*, Wilcoxon rank sum). Moreover, in WT mice L2/3 LFP power was significantly correlated with reaction time on Hit trials (Fig. 5D, linear regression model: 48 ± 12% variance explained within mouse; 18 ± 1% variance explained across mice, *p < 0.05*), whereas L2/3 LFP power in KO mice was less predictive of reaction time (31 ± 8% within mouse; 3 ± 1% across mice, *p = 0.22*). These group differences were not explained by a single underlying relationship between LFP power and reaction times, since aggregating WT and KO data explained even less of the combined RT variance (<1%). Further, elevated 3 – 10 Hz power in KO mice was not significantly correlated with RS neuron firing rates (Fig. 3 – figure supplement 1). Importantly, when we controlled for the effect of contrast on reaction times, (partial correlation), there remained a significant relationship between L2/3 LFP power and reaction times in WT mice, but this relationship was significantly weaker in KO mice (Fig 5D).

Finally, these multiple excitatory activity deficits identified in KO mice performing at perceptual threshold were not the result of differences in stimulus contrast. We re-analyzed neural activity in WT mice at the same contrasts that KO mice required for equivalent performance (Fig. 1K), and found that this did not eliminate differences in stimulus-evoked RS neuron activity, reduced L2/3 LFP responses, or elevated 3 – 10 Hz LFP power (Fig. 5 – figure supplement 1). These results indicate that impairments in both the speed and accuracy of visual perception in KO mice are correlated with and predictable from distinct network-level neural activity deficits in L2/3.

## Discussion

Here we showed that diminished excitatory and elevated inhibitory activity in cortex accompanies impaired sensory perception in a genetically relevant mouse model of ASD. Using a well-controlled, head-fixed visual detection task, we quantified perceptual performance in CNTNAP2^−/−^ KO mice while recording V1 neural activity driven by spatially localized visual stimuli. KO mice detected visual stimuli more slowly and less frequently than WT mice and displayed specific deficits in psychometric sensitivity to contrast. These deficits were accompanied by reduced contrast responses in excitatory neurons and LFP in L2/3, despite threshold contrast being higher in KO mice. Moreover, LFP activity in L2/3 of KO mice was strongly synchronized at low frequencies (3 – 10 Hz) across all trial types. This pervasive low frequency activity in KO mice was detrimental for predicting the speed and accuracy of perceptual performance from L2/3 neural activity. Our results establish that impairments of perceptual performance in a transgenic mouse model of ASD are accompanied by laminar-specific reduction of excitatory activity and abnormal network rhythms.

A first aspect of diminished behavioral performance in KO mice was slower stimulus detection. KO mice took longer than WT mice to detect the same range of stimulus contrasts, and exhibited less improvement of reaction time with higher contrasts; these effects were not explained by lower arousal (assessed via pupil), or by impaired motor vigor (assessed via licking dynamics). Further, delayed behavioral responses were not explained by delayed neural responses in V1. In fact, response latencies in L4 of V1 were nearly identical in KO versus WT mice. This indicates that areas downstream of V1 likely play a role in altered sensory-motor integration that leads to longer reaction times. Although individuals with ASD can also show slower and more variable reaction times (Karalunas et al., 2014), this remains debated (van der Geest et al., 2001; Ferraro, 2016), and the brain areas involved remain unknown. Our study establishes a platform to launch further detailed interrogation of neural activity deficits across multiple brain areas and determine their roles in perceptual impairments and delayed reaction times in transgenic mouse models of ASD.

A second aspect of diminished performance in KO mice was less frequent stimulus detection. Overall, KO mice showed lower Hit rates than WT mice, despite stimuli appearing at the same locations and same visual contrast ranges across all experiments. However, KO mice also showed lower false alarm rates, leading to overall detection sensitivity (d’) that was not statistically different than WT mice. These concerted changes lead to significantly higher response criterion in KO versus WT mice. Importantly, this lower tendency to respond was not explained by lower arousal. Recent studies of visual perception show that both changes in d’ and response criterion contribute to perceptual performance, and that these two components of perception are dissociable with appropriate tasks and likely driven by distinct brain structures (Luo and Maunsell, 2018). Our results here identify that these two aspects of signal detection contribute to overall performance deficits in KO mice, in addition to psychometric deficits in contrast detection, discussed next. Future studies in transgenic mouse models could identify and manipulate brain areas responsible for criterion effects (Sridharan et al., 2017; Crapse et al., 2018; Jin and Glickfeld, 2019) versus those more related to stimulus coding (Goris et al., 2017) to determine contributions of cortical areas to perceptual impairments.

A third aspect of diminished performance in KO mice was reduced psychometric and neural sensitivity to stimulus contrast. First, the stimulus contrast necessary for threshold performance was significantly higher in KO than WT mice, and the rate of rise to saturating performance was significantly shallower in KO mice. Importantly, retinal responses in WT and KO mice were similar across a wide range of stimulus intensities. This degraded relationship between perceptual behavior and stimulus contrast in KO mice was mirrored by weaker contrast dependence of both RS and FS neuron firing during the task. These results establish that reduced V1 neural sensitivity to stimulus contrast accompanies lower perceptual sensitivity to contrast in KO mice. Contrast dependence was markedly reduced in both FS and RS neurons in KO mice during task performance, although FS neurons in awake non-task conditions showed elevated contrast sensitivity. It is conceivable that an “imbalance” of elevated FS activity and reduced RS activity along with a shallower sensitivity to contrast in both FS and RS neurons during behavior contributes to elevated threshold contrast in KO mice.

We revealed that differences in excitatory and inhibitory neuron activity in WT versus KO mice were brain state dependent, and only evident in awake conditions. In awake conditions, visual stimuli evoked overall elevated FS firing and reduced RS firing in KO versus WT mice. This highlights that measurements during anesthesia do not adequately capture cortical neural activity deficits visible in awake conditions. These differences are likely due to the widespread effects of anesthesia on cortical excitation and inhibition (Haider et al., 2013). In awake mice performing the task, reduced RS firing in KO mice was still apparent even though detected stimuli at threshold were static and low contrast, and drove neural activity far less vigorously than drifting, high-contrast stimuli typical of many studies of mouse V1 (Durand et al., 2016; Michaiel et al., 2019). Future studies should consider these factors and compare neural circuit deficits across multiple brain states, stimulus sets, and behavioral outcomes, and also across multiple transgenic mouse models of ASD that display distinct homeostatic adjustments in excitatory and inhibitory activity (Antoine et al., 2019).

Perceptual impairments in KO mice were accompanied by several L2/3 excitatory activity deficits. First, sensory-evoked L2/3 RS firing rates were significantly lower in a manner that depended upon brain state, behavioral outcome, and stimulus contrast. Second, L2/3 LFP responses in KO versus WT mice showed the greatest reduction in contrast sensitivity. Third, L2/3 LFP responses to detected visual stimuli were significantly smaller in KO versus WT mice. Fourth, action potential amplitudes in L2/3 RS neurons in KO mice were significantly smaller than in WT mice. Smaller AP amplitudes in L2/3 RS neurons of KO mice may result directly from loss of CNTNAP2 (a transmembrane protein) that leads to alterations in axonal K^+^ channel localization, deficits in action potential waveforms, and reduced spontaneous synaptic activity in cortical L2/3 pyramidal neurons *in vitro* (Scott et al., 2019), along with deficits in cortical migration of excitatory neurons in L2/3 (Penagarikano et al., 2011). These alterations could in part underlie the activity reductions, smaller extracellular action potentials, and reduced sensory responses measured here in L2/3 during perceptual behavioral impairments. The specific deficits in L2/3 cortical excitatory neurons identified here dovetail with recent findings in humans revealing that individuals with ASD show specific alterations in L2/3 cortical excitatory neurons (Velmeshev et al., 2019). Importantly, L2/3 RS neuron activity provides the major source of feedforward visual information to higher cortical areas (Glickfeld et al., 2013). Recordings of intracellular synaptic and action potentials in L2/3 neurons of awake KO mice could reveal greater insight about the mechanisms underlying these effects and their relationship to sensory and perceptual impairments.

Finally, aberrant 3 – 10 Hz LFP activity in L2/3 of KO mice correlated with perceptual impairments. Several recent studies in WT mice have revealed that 3 – 10 Hz LFP power impairs visual detection (McBride et al., 2019) and predicts perceptual failures (Speed et al., 2019), consistent with the detrimental role of these oscillations in L2/3 for visual coding (Einstein et al., 2017). Here, we found that KO mice showed significantly elevated 3 – 10 Hz power across both correct and incorrect trials; this worsened the predictability of behavioral outcome from LFP activity and also obscured the relationship of LFP activity to reaction times on correct trials. It remains to be seen if 3 – 10 Hz oscillations play a causal role in directly impairing perception in KO mice, or if they are a network-wide consequence of reduced activity in L2/3 excitatory neurons. One recent study directly induced low-frequency oscillations in primate visual cortex, and these caused visual perceptual impairments (Nandy et al., 2019). An intriguing possibility is that low frequency oscillations in L2/3 reduce the dynamic range available to transmit feedforward sensory information to downstream areas during behavior. This suggests that directly attenuating low frequency synchronization in L2/3 neurons may restore excitatory signaling bandwidth and remedy perceptual deficits, a hypothesis that now seems testable in mouse models.

## Methods

### Experimental model and subjects

All procedures were approved by the Institutional Animal Care and Use Committee at the Georgia Institute of Technology and were in agreement with guidelines established by the National Institutes of Health.

#### Surgery

Male C57BL6J (RRID: IMSR_JAX:000664) and CNTNAP2^−/−^ (RRID:IMSR_JAX:017482) mice (5 – 8 weeks old; reverse light cycle individual housing; bred in house) were chronically implanted with a stainless steel headplate with a recording chamber during isoflurane (1-2%) anesthesia. The headplate was affixed to the skull using thin layer of veterinary glue (Vetbond) and secured using dental cement (Metabond). The recording chamber was sealed with a removable polymer (KwikCast). After implant surgery mice were allowed to recover for 3 days before experimentation. During recovery mice were habituated to experimenter handling.

### Behavior

#### Water restriction

Following recovery from surgery, mice were placed under a restricted water schedule (to provide motivation) and trained to detect visual stimuli for water reward. Mice received a daily minimum amount of water (40 ml/kg/day; Burgess et al., 2017; Speed et al., 2019). If mice did not receive their daily minimum water in task, they received supplemental hydration (Hydrogel).

#### Training

Mice first learned to associate visual stimuli with water reward through passive instrumental conditioning. For naïve mice to learn this association, water reward was delivered 0.7s after the onset of a visual stimulus (See “Visual stimuli”, below). Following reward consumption, mice then had to withhold from licking for a mandatory period of time (exponentially distributed intervals from 0.5-6s, randomly selected per trial) in order for visual stimuli to appear on subsequent trials. Lick times were measured with custom built contactless lick detectors (Williams et al., 2018). Typically within 3 – 7 days of training, mice began licking shortly after stimulus onset and prior to reward delivery (anticipatory licking), indicating behavioral responses to the onset of the visual stimulus. Mice were then transitioned to an active paradigm where they only received rewards contingent upon licking during the stimulus presentation (typically 1 s long). On 20% of trials, 0% contrast stimuli were presented in order to measure the probability of licking to the absence of visual stimuli (false alarms). When detection performance was above chance for 2 consecutive days, the contrast and/or size of stimuli were decreased to maintain task difficulty. The main conclusions of this study involve detection of stimuli at a single position in the binocular visual field. Once performance was above chance for a range of low and high contrasts on binocular trials (2 - 33% contrast), we performed acute extracellular recordings.

#### Behavioral metrics

Detection performance was quantified with the psychometric sensitivity index (d’, Green and Swets, 1974), which was calculated as:

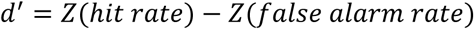

where Z represents the inverse of the normal cumulative distribution (MATLAB function norminv). Response bias or criterion (c) was calculated using the formula:

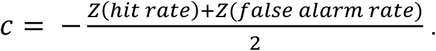

Higher criterion indicates more conservative response bias (withholding responses).

### Recordings

#### Surgical preparation

A small craniotomy (100-400 microns) was opened over binocular V1 during isoflurane anesthesia. Mice were allowed ≥3 hours of recovery before awake acute recordings. There was no difference in behavioral performance in WT mice during recordings (d’: 1.7 ± 0.5) versus the previous day (1.7 ± 0.2, *p = 0.6*, signed rank test). To remove any potential effect of anesthesia or surgery on perceptual performance in KO mice, craniotomies were performed 12-24 hours prior to recordings to ensure equally robust behavioral performance during recordings (d’: 1.7 ± 0.2 versus 1.5 ± 0.3, *p = 0.4*). For both KO and WT anesthetized recordings, mice were given a combination of sedative chlorprothixene (0.1 mg/kg) and isoflurane (0.5-1%), as in our previous studies (Haider et al., 2016).

#### Electrophysiology

Single shank linear 32 site silicon probes (Neuronexus, A1×32) were used to record neural activity across cortical layers. The electrode was typically advanced to 1000 microns below the dura, and the site was covered in sterile artificial cerebrospinal fluid (aCSF). Recordings typically lasted 90 minutes, whereupon the probe was removed and the site cleaned with sterile aCSF and covered with polymer (Kwikcast). Typically we were able to record 3 consecutive days from the same craniotomy.

#### Visual stimuli

During behavior, mice detected Gabor gratings (0.05 - 0.1 cycles/°, σ = 10 - 20°, horizontal orientation, phase randomized per trial). Low contrast (5%) task-irrelevant bars (9° wide, 0.1s duration, inter-stimulus interval of 0.3s, vertical orientation) were also presented during the inter-trial intervals to facilitate receptive field mapping; these faint bars did not affect behavioral performance and they are not analyzed here. After task completion, 100% contrast bars (9° wide, 0.1s duration, inter-stimulus interval of 0.3s, vertical orientation, 100% contrast) were presented across the visual field to map the receptive field. These same bars were used to measure visual responses in awake mice not performing the behavioral task, and also during anesthetized experiments. The bar at the center of the receptive field and the adjacent ±1 bars were used in all subsequent analyses.

#### Eye Tracking

We recorded the animal’s pupil during awake recordings. A high-speed camera (Imaging source DMK 21Bu04.H) with a zoom lens (Navitar 7000) and infrared filter (Mightex, 092/52×0.75) was placed ~22 cm from the animal’s right eye. A near-infrared LED (Mightex, SLS-02008-A) illuminated the eye. Video files were acquired and processed using the Image Acquisition Toolbox in MATLAB with custom code. 1 mm corresponded to ~74 pixels on each frame.

#### Electroretinography

We tested retinal function using full-field flash electroretinography (ERG) as previously described (Mees et al., 2019). Briefly, after overnight dark-adaptation, we anesthetized mice (ketamine 60 mg/kg/, xylazine 7.5 mg/kg) under dim red light, anesthetized corneas with tetracaine (0.5%; Alcon) and dilated pupils with tropicamide (1%; Sandoz). Binocular retinal responses were measured via gold-loop corneal electrodes, with platinum needle electrodes serving as reference and ground in the cheeks and tail, respectively. Testing consisted of 6 scotopic flashes (−4.86 – 2.5 log cd*s/m^2^), followed by 10 minutes of light adaption (30 cd/m^2^) and 3 photopic flashes (−0.2 − 1.4 log cd*s/m^2^). Responses were differentially amplified (1-1500 Hz, 250 ms, 2 kHz) and stored (UTAS BigShot). We measured amplitude and implicit time for a and b waves (Penn and Hagins, 1969) and averaged the traces from right and left eyes for statistical analysis.

### Analysis

#### Spike sorting

Electrical signals were acquired through a Cereplex Direct (Blackrock Microsystems). Raw neural signals were acquired at 30 kHz, and single unit activity was isolated with a semi-automated sorting algorithm (Rossant et al., 2016), as detailed in our previous studies (Speed et al., 2019). We classified single units as fast-spiking (FS, waveform peak-to-trough < 0.57ms) and regular spiking (RS, peak-to-trough > 0.57 ms) based on their waveform widths (Fig. 2-figure supplement 1). FS neurons in mice are predominantly paravalbumin (PV) positive inhibitory neurons, while >85% of RS neurons are putative excitatory neurons (Speed et al., 2019).

#### LFP analysis

Local field potentials were band pass filtered at 0.3-200Hz. Layers were identified via current source density analysis (Niell and Stryker, 2008; Speed et al., 2019) and laminar LFP responses were calculated by taking the average across channels spanning particular layers. We analyzed the residual LFP power in hit and miss trials in the low frequency band (2-20 Hz). We calculated the residual LFP power by fitting the entire power spectrum with a single exponential that excluded the bandwidth of interest. In this bandwidth, residual LFP power is the difference between the measured power and power of the fit, normalized by the fit (Saleem et al., 2017; Speed et al., 2019).

#### LFP ROC Analysis

Receiver Operating Curves (ROC) were constructed to measure the discriminability of hit and miss trial based on low frequency [3-10 Hz] residual power. The area under the receiver operating curve (AUROC) was calculated. Error bars were obtained by bootstrap resampling and repeating the procedure 100 times.

#### LFP-behavior correlations

Reaction times was split into quartiles within each recording. The average residual power or stimulus-evoked spiking activity was then averaged for each quartile. A linear regression model was then fit to the data to determine if there was a correlation between neural activity and reaction time. Error bars were obtained by bootstrap resampling and repeating the fitting procedure 50 times. Distributions of partial correlations controlling for contrasts were constructed using bootstrap resampling (see Fig. 5D).

#### Pupil analysis

Raw video frames were cropped to isolate the eye and pupil. Frames were smoothed with a 2-D Gaussian filter. Based on pixel intensity, the pupil was identified and a least-squares error 2D ellipse was fit to the contours. The pupil area was determined by the amount of pixels in the ellipse. ΔPupil area was calculated as the percent deviation from the mean

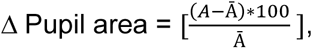

where A is the area in pixels and Ā is the average area across all frames. Similarly, the change in pupil position (azimuth) was calculated by subtracting the average position across all frames.

#### Stimulus-evoked analysis

Visually-evoked firing rates were calculated as the difference between pre-stimulus activity (0.1 s preceding the stimulus onset) and post-stimulus activity (anesthetized: 0 – 0.25 s; awake, no task: 0 – 0.125 s; awake, grating responses during task: 0 − 0.2 s). These windows were chosen based upon the duration of the LFP responses in each condition (Fig. 2 – figure supplement 1). Violin plots of individual data points in all figures show 95% of the data range (±2.5% of range clipped for display). All statistics used full data ranges.

#### Psychometric Analysis

Weibull functions were fit to the hit rate and d’ data (see Fig. 1).

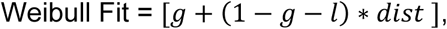

where g is the guess rate, l is the lapse rate, and dist is defined as the cumulative Gaussian distribution:

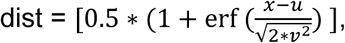

where erf is the error function (‘erf’ in Matlab), u is the mean value, and v, is the standard deviation (Wichmann and Hill 2001).

Stimulus contrast values at Hit rate threshold (0.5) and d’ threshold (1.5) were calculated from fits constructed by bootstrap resampling the data (10% of the data left out, 100 iterations) and generating fits consistent with previous studies (Busse et al., 2011). A d’ threshold of 1.5 was chosen because it falls approximately halfway between the minimum and maximum points of the WT d’ psychometric curve, and is consistent with high-quality perceptual performance as detailed in our previous studies (Speed et al., 2019; Speed et al., 2020).

#### Contrast Sensitivity Analysis

Stimulus evoked single unit activity curves for passive awake recordings were constructed from responses to flashing bars at the center of the receptive field (5-100% contrast). Single unit activity during behavior was binned between 5-35% (bin size = 10; see Fig. 3). A linear regression model was fit to the firing rate data as a function of contrast to calculate the slope. A bootstrap resampling approach as described above (100 resamples) was used to construct distributions of estimated slopes.

#### Awake LFP Visual Responses Analysis

Weibull functions were fit to the LFP contrast response curves (see Fig. 4). A resampling approach as described above was used to construct distributions of the difference in LFP amplitude between WT and KO mice.

### Experimental design and statistical analysis

Our experimental design centered on measuring neural activity and behavior during identical sensory conditions, and comparing these across KO and WT mice matched in age, sex, recording region, and methods. Experimenters were not blinded to the group identity of each subject, but this was not required for performing identical measurements of behavioral performance or electrophysiology. We performed these studies in comparable numbers of subjects and experiments across groups. Throughout this paper, unpaired comparisons utilized Wilcoxon rank sum tests (tails specified) or sign tests (for differences from scalar values), and paired comparisons utilized Wilcoxon signed rank tests, unless otherwise noted.

### Data availability

All data structures and code that generated each figure are available upon reasonable request.

## Author contributions

J.D.R., H.A., A.S. performed behavioral experiments; J.D.R., A.S. performed electrophysiological experiments; C.M. performed ERG experiments and data analysis with M.P.; J.D.R., A.S., B.H. performed data analysis; J.D.R. and B.H. wrote the manuscript with feedback from all authors.

## Acknowledgements

J.D.R. was funded by a Goizueta Foundation fellowship and Alfred P. Sloan Foundation’s Minority Ph.D. (MPHD) Program Fellowship. B.H. was funded by the GT Neural Engineering center (1241384), the Whitehall Foundation, the Sloan Foundation, NIH NINDS (NS107968), NIH BRAIN Initiative (NS109978), and the Simons Foundation Autism Research Initiative (SFARI). M.P. was funded by Department of Veterans Affairs Rehabilitation Research and Development Service Merit Award (I01RX002615) and Senior Research Career Scientist Award (IK6RX003134).

---

**Figure 1–figure supplement 1.**
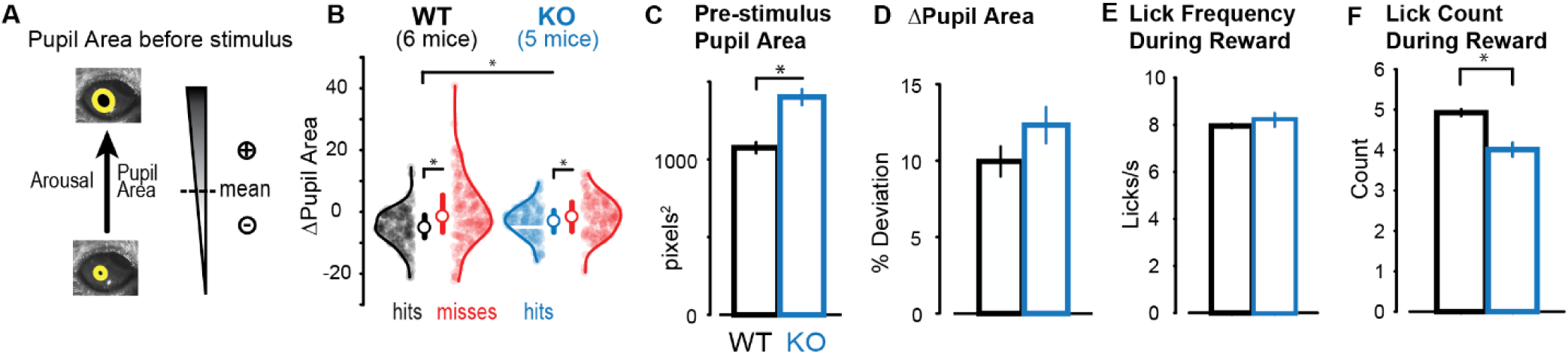
Arousal and motor activity do not explain perceptual impairments. **A.**Pupil area was measured during each trial. On a given trial, larger positive deviation of pupil area from session mean indicates higher arousal. **B.** ΔPupil area preceding stimulus onset on Hit trials was significantly smaller than Miss trials in both WT (Hits: −5.2 ± 0.5%; Misses: −0.5 ± 0.8%; n = 190 sessions in 6 mice; mean ± SEM throughout the figure; *p< 0.01*, rank sum test) and KO mice (Hits: −3.2 ± 0.5%; Misses: −1.5 ± 0.6%; n = 138 sessions in 5 mice; *p<0.01*, rank sum test). ΔPupil area preceding Hits was significantly smaller in WT versus KO mice (*p< 0.01*, rank sum test), but not for Misses (*p=0.65*, rank sum test). ΔPupil area was calculated as the frame-by-frame percent deviation from the mean pupil area of the whole recording session (see Methods). Median ± IQR is plotted inside the distributions. White horizontal line in KO Hits distribution indicates mean of WT Hits. **C.** Overall mean pupil area was larger in KO versus WT mice (WT: 1075 ± 35 pixels^2^, KO = 1402 ± 51 pixels^2^; *p<0.01*, rank sum test). **D.** ΔPupil area during reward consumption was not significantly different between WT (10.0 ± 1.0%) and KO mice (12.3 ± 1.2%; *p=0.57*, rank sum test). **E.** Lick frequency during reward was not significantly different between WT (8.0 ± 0.1 licks/s) and KO mice (8.2 ± 0.3; *p=0.12*, rank sum test). **F.** Lick count during reward bouts was significantly smaller (by one lick) in KO mice (4.0 ± 0.2) compared to WT mice (5.0 ± 0.1, *p < 0.01*, rank sum test).

**Figure 1–figure supplement 2.**
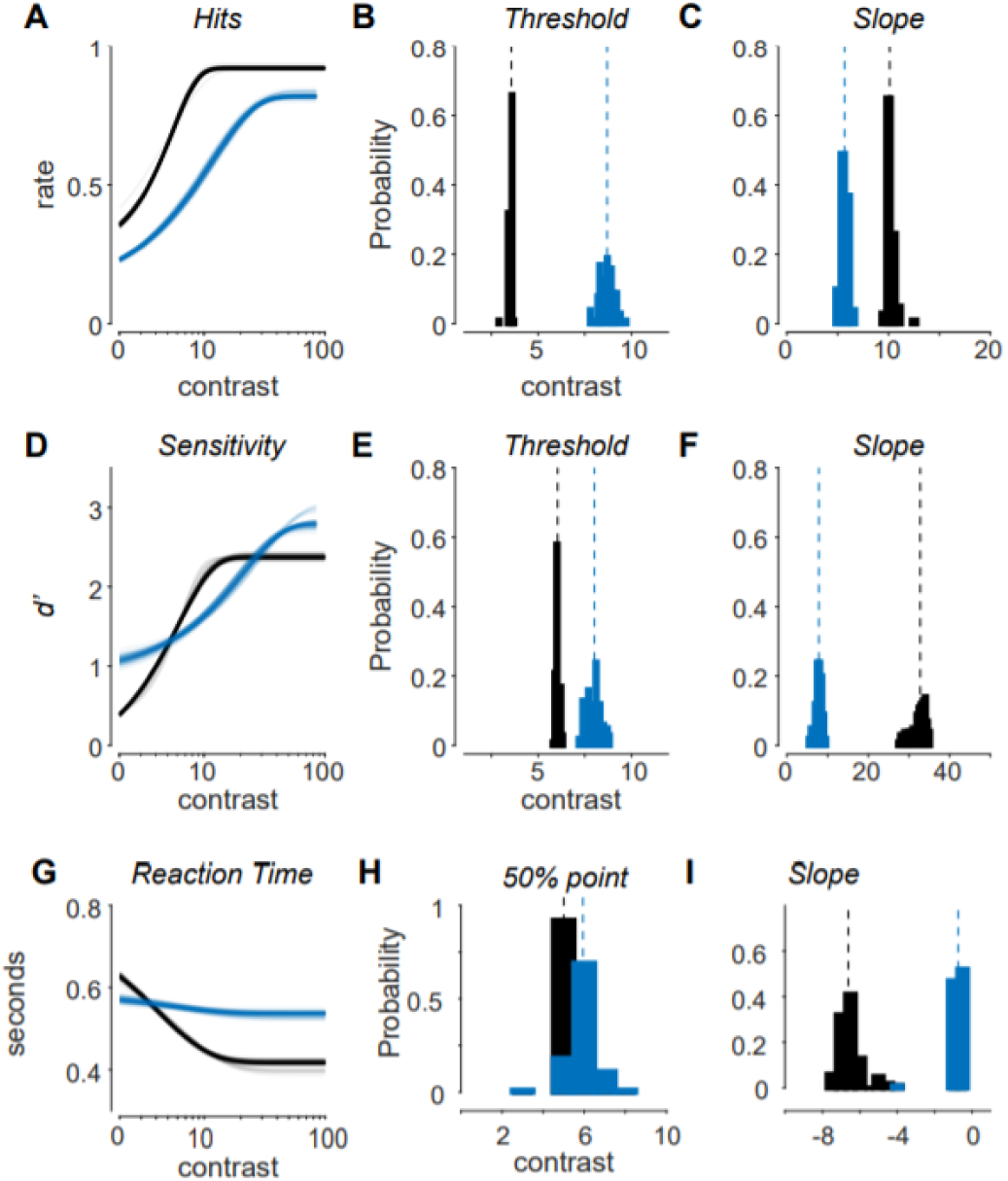
Psychometric performance differs between WT and KO mice. **A.** Psychometric curve fits in WT and KO mice. Each curve was fit from bootstrap resampling (random 10% left out; WT: 19,288 Hit trials in 187 behavioral sessions; KO: 6,501 Hit trials in 71 behavioral sessions). 100 overlaid curves fit to resampled data (throughout figure). **B.** Distribution of psychometric discrimination threshold contrasts (at 0.5 Hit rate) is significantly greater in KO (8.7 ± 0.4% contrast, mean ± SD, throughout figure) versus WT mice (3.5 ± 0.1% contrast, *p<0.01*, Wilcoxon rank sum test). Threshold contrasts from each fit in A, means at dashed line. **C.** Distribution of maximum slope of hit rate psychometric curves is significantly shallower in KO (5.7±0.3) versus WT mice (10.2 ± 0.4, *p<0.01*, Wilcoxon rank sum test). **D.** Same as A, for d’. Same resampling as A for curve fits. **E.** Same as B, for d’. Discrimination threshold contrasts (defined by d’ = 1.5) significantly greater in KO mice (8.0 ± 0.4% contrast) versus WT mice (6.0 ± 0.1% contrast, *p<0.01*, Wilcoxon rank sum test). **F.** Same as C, for d’. Distribution of d’ psychometric curve slope significantly shallower in KO (7.8 ± 0.8) versus WT (32.2 ± 1.9, *p<0.01*, Wilcoxon rank sum test) **G.** Exponential fits of reaction time (RT) as a function of contrast. Same resampling as in A-F. **H.** Contrast level at which RT improved by 50% is significantly higher in KO than WT mice (WT: 5.1 ± 0.3% contrast; KO: 5.9 ± 0.7%, *p<0.01*). **I.** KO mice have a shallower (more positive) maximum slope of RT improvement with contrast (WT: - 6.5 ± 0.6; KO: −0.8 ± 0.4, *p<0.01*). Slopes negative since RT decreases with contrast.

**Figure 2–figure supplement 1.**
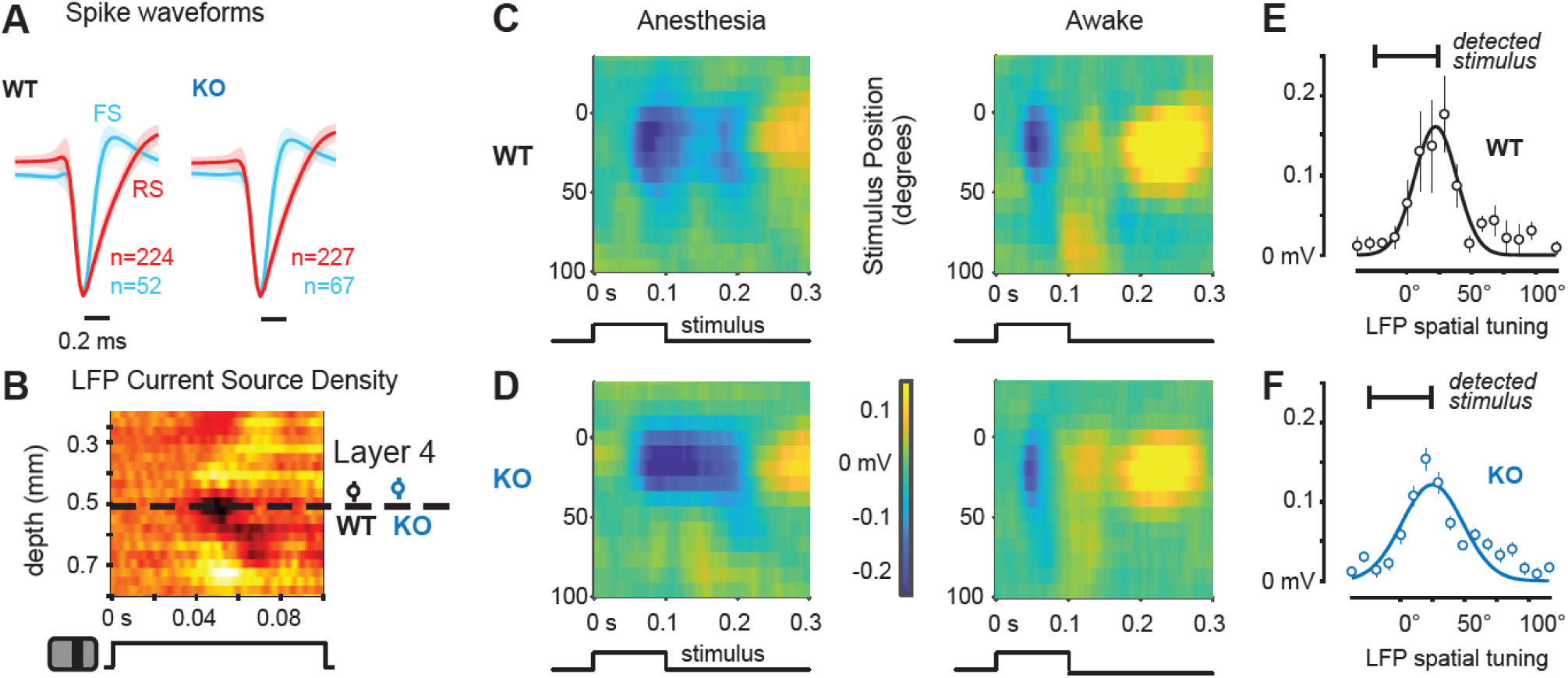
Cell types, laminar estimates, receptive fields, and stimulus positioning were similar across WT and KO recordings. **A.** Population spike waveforms of fast-spiking (FS) putative inhibitory neurons and regular spiking (RS) putative excitatory neurons in WT and KO mice (mean ± SEM). Individual waveforms normalized to maximum negative deflection before averaging. **B.** Left, Example current source density (CSD) heatmap showing response to high contrast bar onset in best spatial location. Rows indicate LFP sampled across cortical layers, stimulus time course at bottom. Dark colors identify earliest and largest current sink, presumed to be layer 4 (L4). Right, population estimates of L4 from CSD are similar across WT (0.51 ± 0.02 mm, mean ± SEM, n = 23) and KO mice (0.51 ± 0.02 mm, n = 28). **C.** Spatial tuning of local field potential (LFP) responses in V1 of WT mice recorded during anesthesia (left; n = 13 recordings, 4 mice) and wakefulness (right; n = 10 recordings, 4 mice). Ordinate indicates stimulus (bar) position (in azimuth, vertical meridian at 0°), abscissa shows time course. Note that both WT and KO spatial tuning peaks within the central 20° of the visual field, in the binocular zone. **D.** Same as c, for KO mice during anesthesia (left, n=18 recordings, 4 mice) or wakefulness (right, n = 10 recordings, 4 mice). Note that state-dependent amplitude and time course of LFP spatial tuning is similar across KO and WT recordings. **E.** Spatial tuning of population LFP response in awake WT mice (line shows Gaussian fit, peak at 22°). The position and extent of stimuli detected during the task shown at top. **F.** Same as E, for KO mice. Gaussian fit peaks at 24°. Receptive field size (half-width of peak-normalized Gaussian fit) was not significantly different between WT (18 ± 5°, mean ± SD) and KO (15 ± 4°) mice during anesthesia (*p=0.29*, rank sum test) or wakefulness (19 ± 14°; 18 ± 9°; *p=0.96*, rank sum test).

**Figure 2–figure supplement 2.**
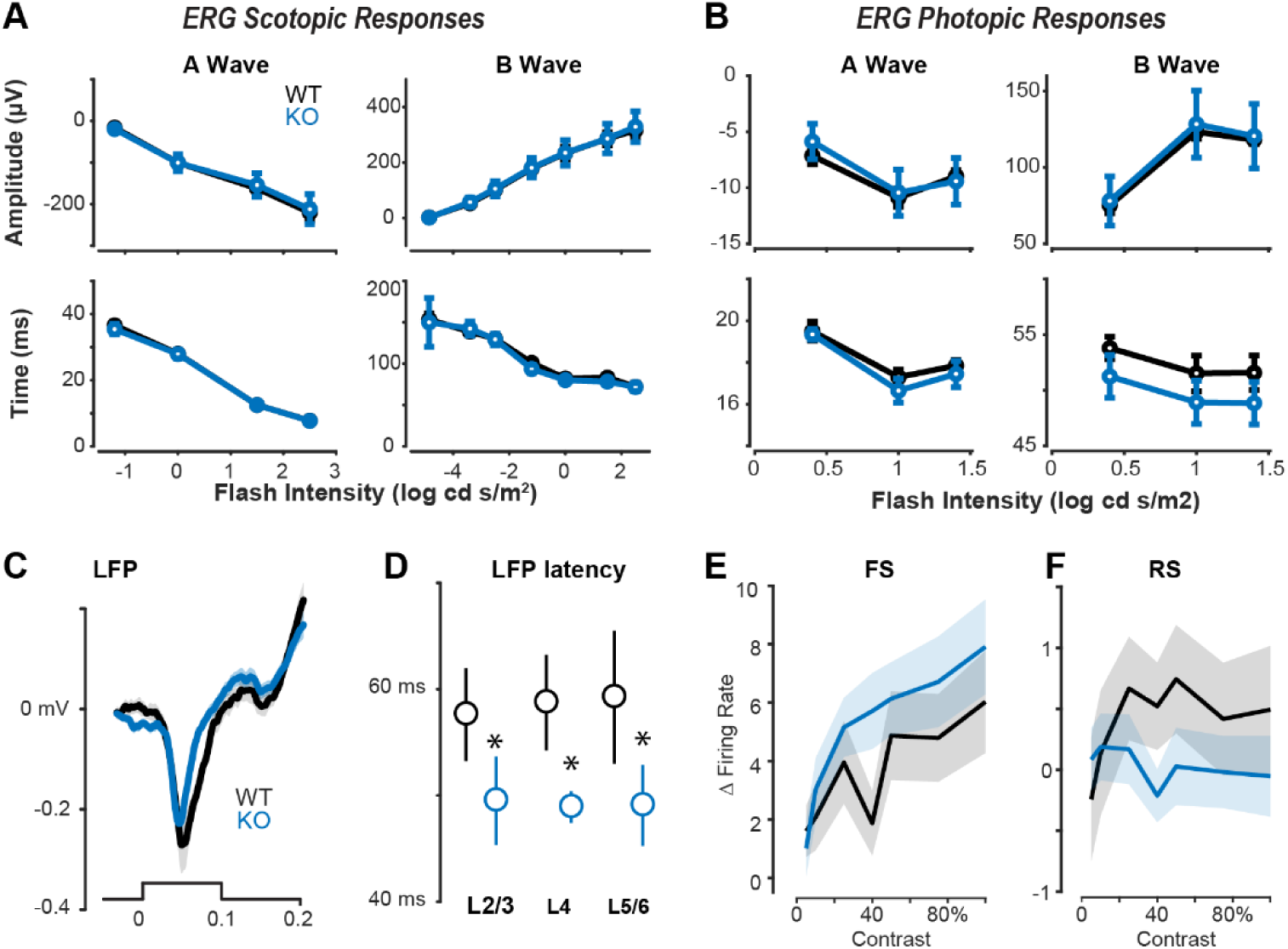
Retinal responses, LFP latencies, and contrast responses in WT and KO mice. **A.** Electroretinography (ERG) of scotopic responses (dark adapted; rod-dominated) as measured by the peak amplitude (top rows) and implicit time of the peak amplitude (bottom rows) for the initial negative deflection (a Wave, left) and the subsequent positive deflection (b Wave, right), which corresponds to the hyperpolarization of photoreceptor cells and depolarization of bipolar cells (Penn and Hagins, 1969). Neither the peak amplitude (a wave: *p=0.94*; b wave: *p=0.94*, n = 4 WT and 4 KO mice) nor implicit time (a wave: *p=1*; b wave: *p=0.52*) were significantly different between WT and KO mice. Mean ± SEM is plotted throughout the figure. See (Mees et al., 2019) for detailed description of ERG Methods. **B.** Same as a, but for photopic responses (light adapted; cone-isolating). Neither the peak amplitude (a wave: *p=0.99*; b wave: *p=0.58*, 4 WT and 4 KO mice) nor implicit time (a wave: *p=0.62*; b wave: *p=0.08*) were significantly different between WT and KO mice. **C.** Population average LFP response across all layers to best bar (100% contrast, center of receptive field) for awake WT and KO mice. Mean ± SEM. **D.** LFP response latency was significantly faster in KO mice across all layers. L2/3: WT = 58 ± 42 ms, KO = 50 ± 4 ms (*p=0.05*, rank sum test); L4: WT = 59 ± 4 ms, KO = 49 ± 1 ms (*p=0.01*, rank sum test); L5/6: WT = 59 ± 6 ms, KO = 49 ± 4 ms (*p<0.01*, rank sum test). **E.** Spiking responses of FS neurons as a function of contrast to the single best bar (9° wide) presented at the center of the receptive field. KO mice had a steeper contrast dependence than WT mice (linear fits, slope of firing rate versus contrast: KO, 6.1 ± 0.3; WT, 4.3 ± 0.3, *p<0.01*, Wilcoxon rank sum, mean ± SD). **F.** Same as E for RS neurons. KO mice had weaker contrast dependence than WT mice (linear fits, slope of firing rate versus contrast: KO, 0.5 ± 0.05; WT, −0.21 ± 0.02, *p < 0.01*, Wilcoxon rank sum, mean ± SD).

**Figure 3–figure supplement 1.**
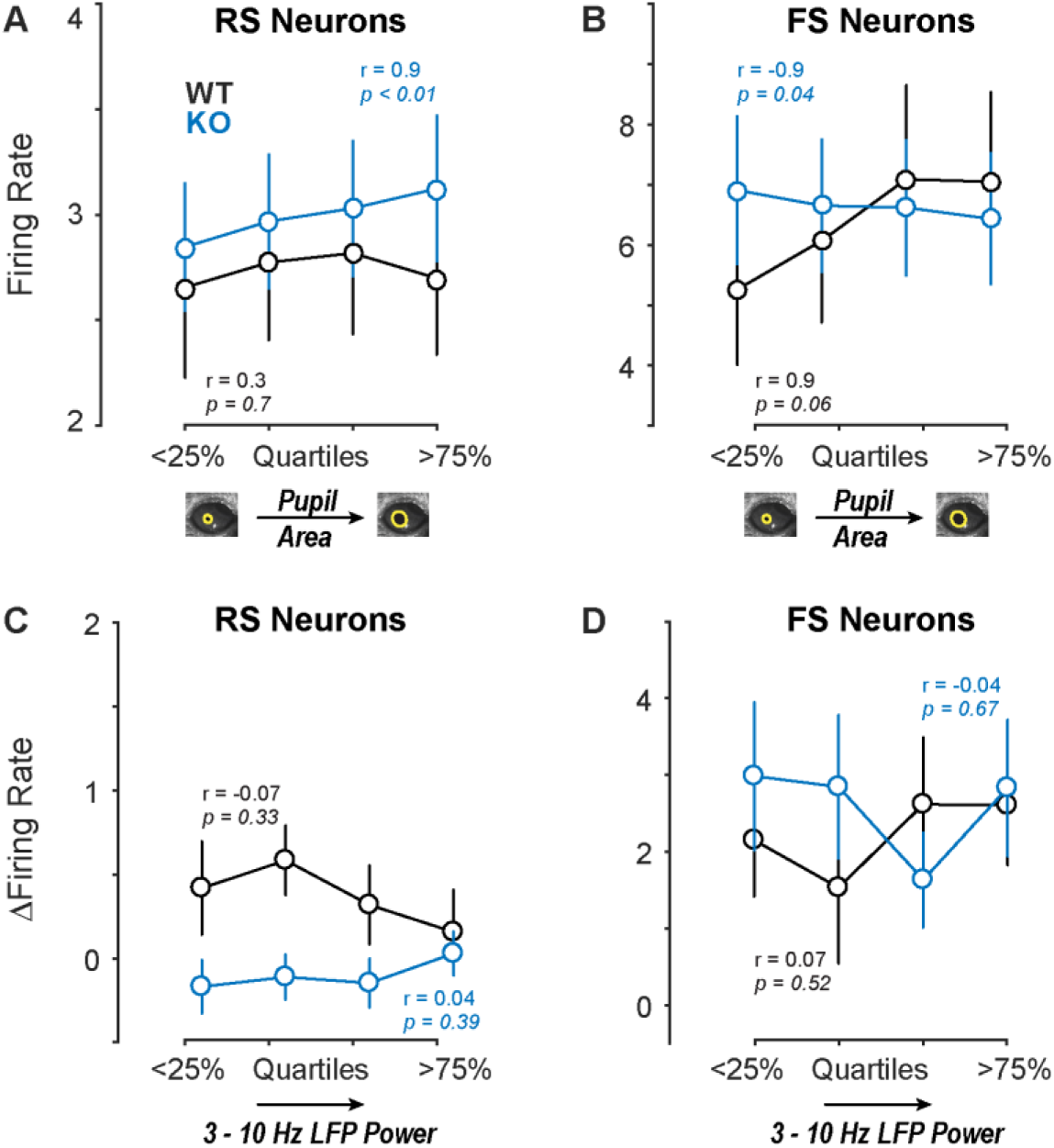
Firing rates in WT and KO mice as a function of pupil area and low frequency LFP power during behavioral task. **A.** RS neuron firing rate preceding stimulus onset, sorted by single-trial pupil area quartile (relative to mean area overall during session mean area). Significant positive correlation of RS firing rate and pupil area in KO mice (r = 0.98, *p < 0.01*, n = 103 neurons, 1677 trials, 15 recordings in 3 mice) but not WT mice (r = 0.31, *p = 0.68*, n = 49 neurons, 744 trials, 7 recordings in 3 mice). **B.** Same as a, for FS neuron firing. Significant negative correlation of FS firing rate and pupil area in KO mice (r = −0.94, *p = 0.03*, n = 38 neurons, 1677 trials, 15 recordings in 3 mice) but not WT (r = 0.96, *p = 0.06*, n = 20, 744 trials, 7 recordings in 3 mice). **C.** Stimulus-evoked RS neuron firing rate (above baseline), sorted by 3 — 10 Hz LFP power quartiles. No significant correlation in KO (r = 0.04, *p = 0.39*, n = 103 neurons, 1677 trials, 15 recordings in 3 mice) or WT mice (r = −0.07, *p = 0.34*, n = 20, 744 trials, 7 recordings in 3 mice). **D.** Same as c, for FS neuron firing. No significant correlation in KO (r = −0.04, p = 0.67, n = 103 neurons, 1677 trials, 15 recordings in 3 mice) or WT mice (r = 0.07, *p = 0.52*, n = 20, 744 trials, 7 recordings in 3 mice).

**Figure 3–figure supplement 2.**
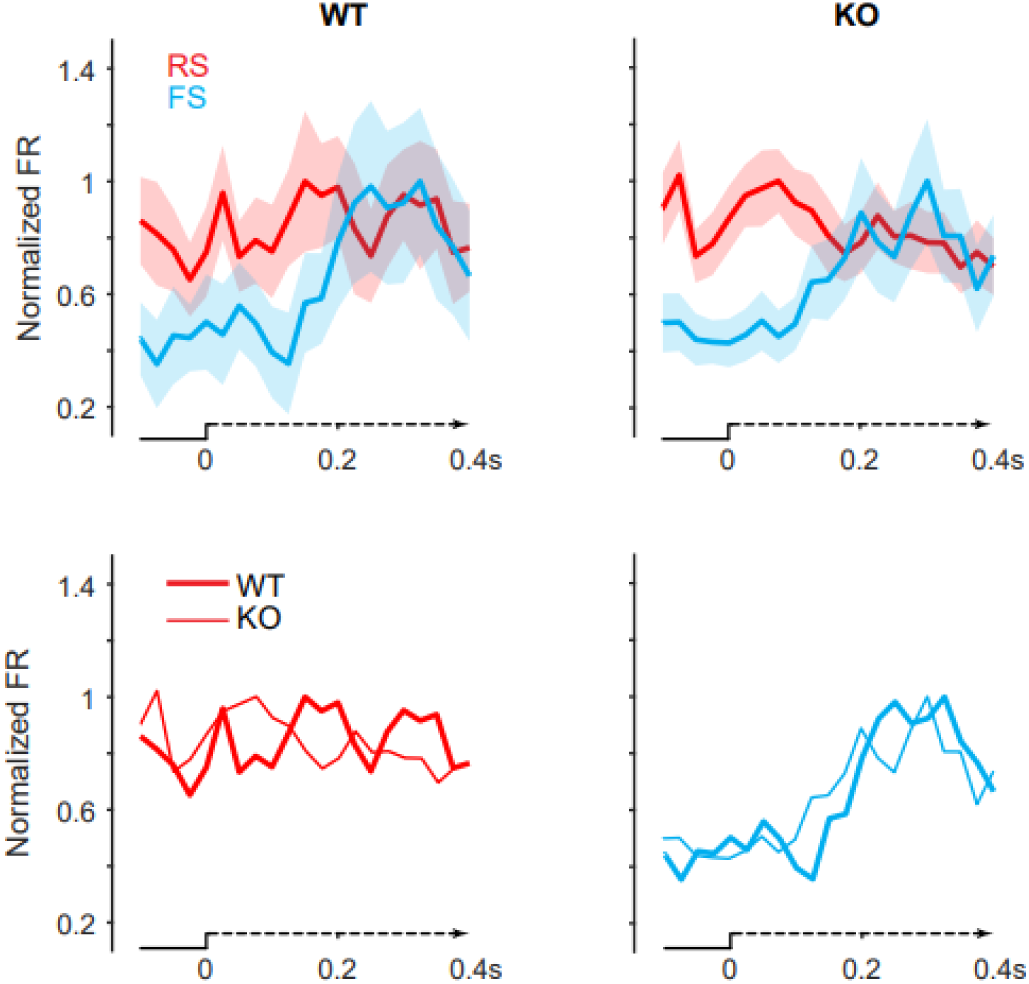
RS and FS responses on correct detection trials. Top Row, Normalized PSTHs of RS and FS neurons in WT mice (left) and KO mice (right) during perceptual detection. Bottom Row, Normalized RS (left) and normalized FS (right) PSTHs of WT and KO mice during perceptual detection.

**Figure 3–figure supplement 3.**
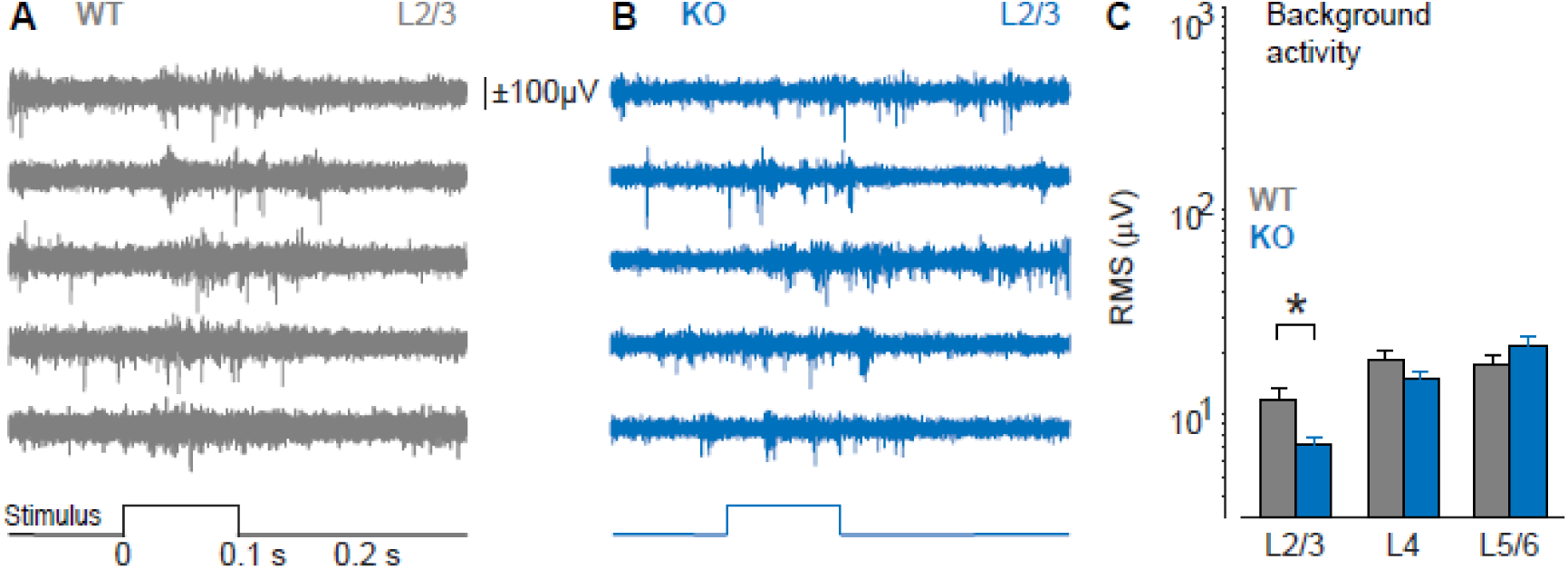
Background activity and spiking in WT and KO recordings. **A.** Example single-trial raw signal (high pass >300Hz) from L2/3 in an awake WT mouse. 5 adjacent channels (25 micron spacing) show clear stimulus evoked spiking (bottom, 100% contrast flashed bar presented outside of behavioral task). **B.** Same as a, for L2/3 in awake KO mouse. Note that unresolved background activity (“hash”) is not larger in KO recording versus WT recording. **C.** Root mean square (RMS) amplitude of background activity in L2/3 of KO mice is smaller than in WT mice (7 ± 1 μV vs 12 ± 2 μV, *p = 0.01*). No significant difference in other layers (L4: 15 ± 2 μV vs 19 ± 2 μV, *p = 0.14*; L5/6: 22 ± 2 μV vs 18 ± 2 μV, *p = 0.28*, rank sum test).

**Figure 5–figure supplement 1.**
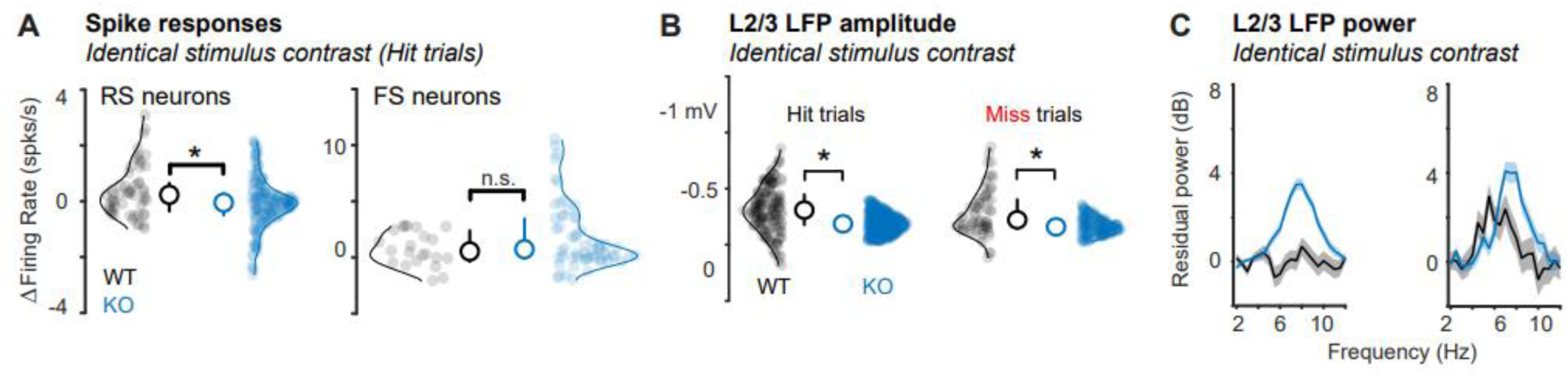
Reduced RS firing, smaller LFP responses, and elevated 3 – 10 Hz LFP activity of KO mice is not due to higher stimulus contrast. **A.** Spiking responses of RS neurons (left) and FS neurons (right) during hit trials for the same contrasts (5-15%). RS neuron responses in KO mice are significantly reduced (WT = 0.47 ± 0.20 spikes / s, 49 neurons, mean ± SEM throughout the figure; KO = −0.07 ± 0.11 spikes / s, 103 neurons, *p<0.05*, 1-tail Wilcoxon rank sum throughout the figure). There was no significant difference in FS activity (WT = 0.87 ± 0.43 spikes / s, 20 neurons; KO = 2.25 ± 0.59 spikes / s, 38 neurons *p=0.21)*. Median ± IQR plotted. **B.** For identical contrasts, L2/3 LFP amplitudes were significantly reduced in KO mice on hit trials (Left: WT = −0.32 ± 0.03 mV, 160 trials; KO = −0.19 ± 0.004, 588 trials, *p<0.01*) and miss trials (Right: WT = −0.28 ± 0.04 mV, 67 trials; KO = −0.17 ± 0.006 mV, 188 trials, *p<0.01*). **C.** For identical contrasts, integrated L2/3 LFP power (3-10Hz) was elevated in KO mice on hit trials (Left: WT = 0.24 ± 1.58, 160 trials; KO = 12.18 ± 1.01, 588 trials, *p<0.01*) and miss trials (Right: WT = 7.59 ± 2.11, 160 trials; KO = 12.41 ± 1.37, 588 trials, *p<0.05*). Integrated 3-10Hz residual power was greater on miss vs. hit trials in WT mice (*p<0.02*), but not in KO mice (*p=0.45*).

